# COVID-19 genetic risk and Neanderthals: A case study highlighting the importance of scrutinizing diversity

**DOI:** 10.1101/2020.11.02.365551

**Authors:** Inken Wohlers, Verónica Calonga-Solís, Jan-Niklas Jobst, Hauke Busch

## Abstract

Recent genome wide association studies (GWAS) have identified genetic risk factors for developing severe COVID-19 symptoms. The first published study reported a 1bp insertion rs11385942 on chromosome 3 (1) and subsequent studies single nucleotide variants (SNVs) such as rs35044562, rs67959919 (2) and rs13078854 (3), all highly correlated with each other. Zeberg and Pääbo (4) subsequently traced them back to Neanderthal origin. They found that a 49.4 kb genomic region including the risk allele of rs35044562 is inherited from Neanderthals of Vindija in Croatia. Here we add a differently focused evaluation of this major genetic risk factor to these recent analyses. We show that (i) COVID-19-related genetic factors of three previously assessed Neanderthals deviate from those of modern humans and that (ii) they differ among world-wide human populations, which compromises risk prediction in non-Europeans. Currently, caution is thus advised in the genetic risk assessment of non-Europeans during this world-wide COVID-19 pandemic.

## Main

In general, GWAS relate genotypes to phenotypes such as disease susceptibility and severity. However, association does not imply causality. To pinpoint causal variant(s) underlying a GWAS association signal which typically comprises many correlated variants, a so-called fine-mapping is performed in a first step. And ultimately, fine-mapping must be followed by experimental validation to eventually identify causal variant(s) and mechanisms. While GWAS are based on cohort data, a personal risk can be assessed nonetheless, via associated variants as proxies for causal variants. For this, the cohort’s genetic linkage patterns need to be representative of the individuum’s genetic background. This requirement, however, is often violated, especially for individuals having non-European ancestry. In a world-wide COVID-19 pandemic this might jeopardize individual genetic risk prediction and requires current risk factors to be used with caution as we show below.

The first and to date only GWAS published in a peer-reviewed journal by the Severe Covid-19 GWAS Group *et al*. (1) obtained a credible set of 22 highly correlated risk variants through *in silico* fine-mapping using FINEMAP (5), including the reported lead variant rs11385942. In combination, these variants have an overall probability greater than 95% to include the causal variant, while each of the 22 variants has an individual causal probability between 1 and 11% (median: 4%). This initial study was a Meta-Analysis of a Spanish and an Italian cohort (overall n=1610 cases). The COVID-19 host genetics initiative included this data into a world-wide Meta-GWAS currently in release 4 (n=8638 cases). Fine-mapping of this latest release results in a credible set of only 10 variants, which are a subset of the published 22 credible set variants. This fine-mapping does not utilize linkage disequilibrium information of the GWAS cohort itself, which is the gold-standard setting for fine-mapping and which has been applied to the initial Spanish and Italian cohort. Furthermore, it may be affected by differences between European and non-European cohorts, for example if a causal variant does not occur in all cohorts. World-wide genetic diversity of risk haplotypes identified within a relatively homogeneous GWAS cohort should thus be scrutinized to help with the interpretation of findings and to identify early possible limitations, as we do exemplarily in the following.

19 of these 22 risk variants are identifiable in the Vindija Neanderthal genome and four carry protective alleles, which accumulates to a risk probability of 64% for containing a causal variant. This probability could increase to 82%, if the missing variants were risk alleles as well. Zeberg and Pääbo addressed 13 of the 19 variants in their Neanderthal study, with an overall maximum probability to include a causal variant of 61%. However, as two positions carry protective alleles the risk probability of the previously assessed Vindija Neanderthal haplotype is only 52%.

Our results presented here complement the haplotype-based assessment of Zeberg and Pääbo. We use the same 1000 Genomes (6) data as in the original study, but with three important differences: (i) We investigate haplotypes within a larger genomic region of 65.8 kb length that incorporates all 22 COVID-19 related variants, all of which have an overall probability of more than 95% to include the causal variant. This considerably increases the probability of only 61% covered by the original analysis. (ii) We investigate only the haplotypes for the 22 credible set variant positions, that is, only COVID-19 risk-related haplotypes. Previously, all haplotypes including all variant positions were used to obtain a comprehensive phylogenetic tree of the locus, which showed how haplotypes carrying the latest lead variant rs35044562 form a clade with Neanderthals. Here we characterize risk-related haplotypes irrespective of phylogenetic relationships. (iii) Lastly, the former haplotype-based assessment used only lead variant rs35044562 to classify haplotypes as risk ones. Instead, we here make use of individual probabilities of the risk variants.

Haplotypes from 1000 Genomes belong to 38 different haplogroups (labeled H1-H38 in order of overall count, Fig. 1c). Risk-related haplotypes have an aggregated frequency of 10% in the whole dataset and variable frequencies from 1 to 31% in different continental populations (Fig. 1a). Eight haplogroups, H1-H8, have counts higher than 10 and the most common is the protective haplogroup H1. Risk haplotypes of groups H2-H8 tend to differ between continental populations (Fig. 1a). For them, COVID-19 genetic risk probability varies substantially between 8 and 96%. The high risk haplogroups H2, H3 and H8 differ by one or two alleles, and differ from the low risk haplogroups H5, H6 and H7 all of which are similar to the protective haplogroup H1 (Fig. 1b). However, individuals carrying a risk haplogroup very dissimilar from Neanderthal haplotypes may still carry a causal variant (Fig. 1c); this holds particularly true for Africans with haplogroups H5 or H6 (19% or 11% probability) and for Asians with haplogroup H7 (8% probability). Haplogroup H3 has highest risk probability and is the most common risk haplogroup in Europeans and Americans (Fig. 1b).

**Figure 1:**
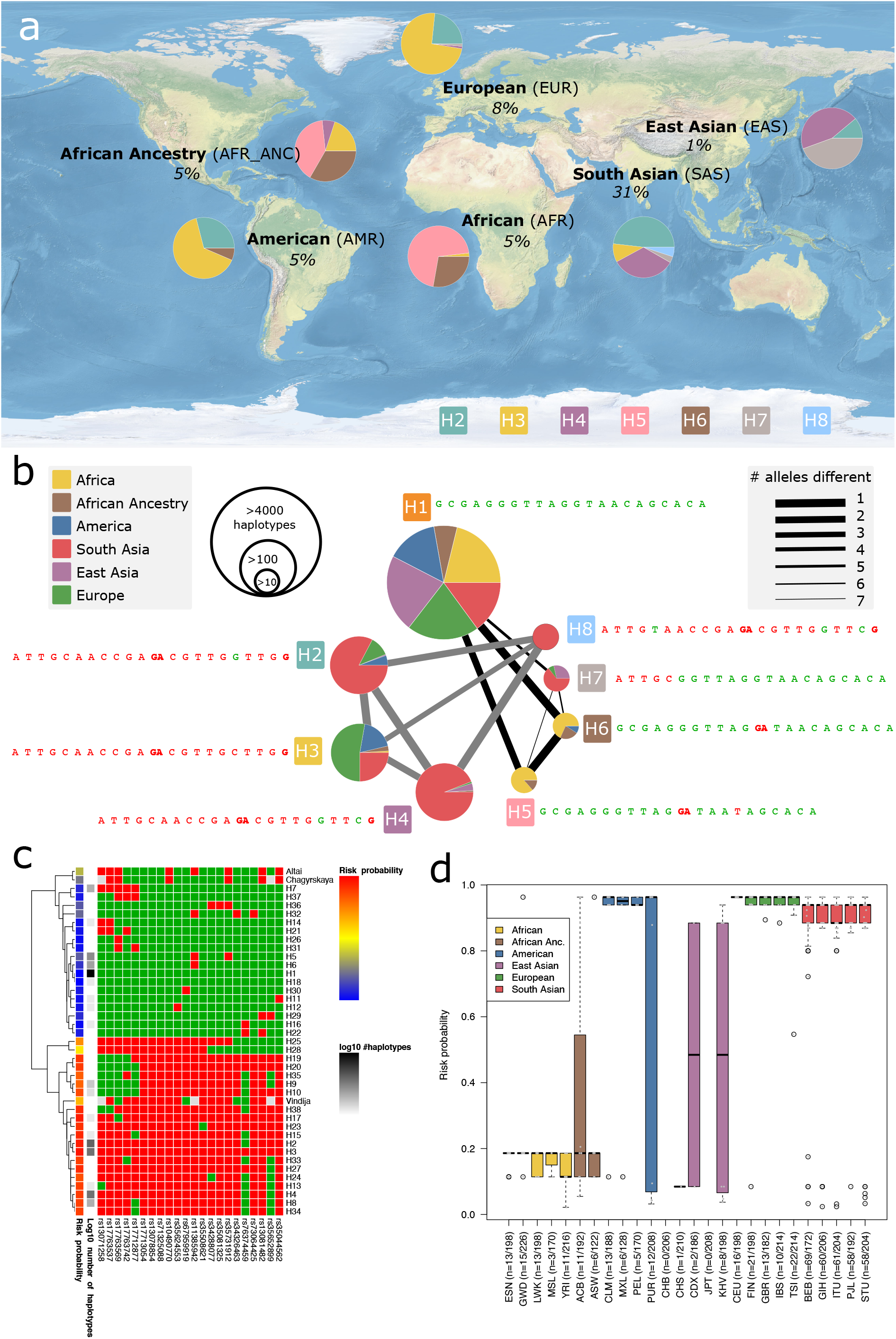
a) Risk haplogroups in different continental populations. b) Comparison of the eight most common haplogroups and their occurrence in continental populations. For each haplogroup, the alleles for 22 risk variants are provided in the order of chromosomal position. Red alleles are risk alleles, green alleles are protective. Network nodes and edges are correlated with haplotype frequency and allele difference, respectively. c) Heatmap depicting all 38 haplogroups observed in n=5008 haplotypes from 1000 Genomes as well as three Neanderthals (Altai, Chagyrskaya and Vindija). Red denotes risk alleles, green denotes protective alleles. Variant order is according to chromosomal position. Annotated are the number of haplotypes of every haplogroup (log10 scale) and the risk probability of every haplogroup considering 22 fine-mapped variants. d) Distribution of risk probabilities for risk haplotypes of the 1000 Genomes populations. Box plots display median and lower/upper quartiles; whiskers denote the most extreme data point no more than 1.5 times the interquartile range; outliers are data points extending beyond whiskers.

All human risk haplogroups differ from the three previously assessed Neanderthal haplotypes (Fig. 1c). They share at most 11 of the 13 previously assessed Neanderthal alleles and 16 of 19 known Vindija alleles. We used IBDmix (7), a recent tool for individual-level identification of Neanderthal-inherited regions, to obtain Vindija Neanderthal-introgressed sequences greater than 30 kb. Introgressed sequences overlapped the considered 65.8 kb genomic region for most risk haplogroup carriers H2, H3, H4 and H8, yet were absent for most protective homozygous H1 haplogroup carriers and for all low risk H5/H6 haplogroup carriers, respectively. Thus, because all high risk haplogroups are located within Neanderthal introgressed region, the lack of 3 of 19 risk-associated alleles shared by all human high risk haplogroups is best explained by genetic diversity within Neanderthals – the introgressed sequence seems to originate from a different Vindija Neanderthal than the one assessed.

In Africans, the protective H1 and low risk H5/H6 haplogroups occur almost exclusively. Still this population carries the lead risk variant allele rs11385942 as well as, interestingly, two protective Neanderthal alleles. Given the only 11% probability of rs11385942 to be causal there is thus a fair chance that this lead variant incorrectly classifies Africans to be at risk of developing severe COVID-19 symptoms. This would contradict classification using the lead variant rs35044562. Overall, when classifying individuals that carry the GA allele of rs11385942 to be at risk, 477 haplotypes would be considered at risk, and these have an average probability of only 82% to contain the causal risk allele. If instead the Meta-GWAS risk allele of rs35044562 were used for classification, African haplogroups H5 and H6 would not be considered at risk. The overall 410 haplotypes considered at risk have an average probability of 92% to contain the causal risk allele.

Only 1% of East Asian haplotypes belong to the risk haplogroups and none of them belongs to the largest risk probability haplogroup H3, which is predominantly European. Contrary to this, the South Asian risk haplotype frequency is 31%, the highest among all continental populations, a consequence of the predominance of the haplogroups H2, H4 and H8. These haplogroups contain protective alleles that reduce the risk probability by 2, 8 and 9% with respect to the highest risk haplogroup H3. Most South Asian haplotypes thus have lower risk probability than European haplotypes. Zeberg and Pääbo denoted the difference between South and East Asian populations as unexpected and significant and state that it may indicate genetic selection. Our analysis shows that South Asian risk haplogroups are genetically more diverse, which may be the result of adaption. Both East Asian risk haplotype depletion as well as South Asian haplotype diversity can be hypothesized to result from exposure to pathogens related to severe respiratory diseases. Further, the protective G allele of rs76374459 is shared by predominantly South Asian haplogroups H2, H4, H7 and H8. If this variant was causal (2% probability) using lead variants such as rs11385942 or rs35044562 would incorrectly classify individuals carrying these haplogroups to be at risk. This applies to few Europeans, but mostly to non-Europeans.

In conclusion we find that classification into high and low COVID-19 risk is error-prone in non-European populations, if this assessment is based on European risk variants and probabilities, especially when using lead variant rs11385942, which is the only one to date published in a peer-reviewed journal. The risk haplogroup diversity observed across populations thus compromises risk assessment in non-Europeans. This situation is currently improved by world-wide GWAS efforts also in non-European populations, by Meta-GWAS, as well as by trans-ethnic GWAS. With respect to the latter, a recent GWAS performed by the company 23andMe replicates the major genetic risk factor addressed here, has rs13078854 as lead variant and a credible set of 20 variants (3). Further, in-silico fine-mapping for the latest release of the Covid19 Host Genetics Initiative results in only 10 credible set variants, all of them distinguishing common high risk haplogroups from common low risk haplogroups – these are 10 of 13 variants used in the Neanderthal study. Thus, various genetic studies already pinpoint, quantify and limit the set of candidate causal variants, and a combined population genetic view will help narrowing down the list using in silico as well as complementary, e.g. experimental approaches. These diverse systems genetics efforts will eventually converge into genetic causes and corresponding molecular mechanisms that explain non-environmental variation in COVID-19 severity.

## Acknowledgements

We thank the COVID-19 Host Genetics Initiative for publicly releasing GWAS summary statistics. IW and HB acknowledge funding by the Deutsche Forschungsgemeinschaft (DFG, German Research Foundation) under Germany’s Excellence Strategy—EXC 22167-390884018. Verónica Calonga-Solís was supported by a scholarship from Deutscher Akademischer Austauschdienst (DAAD) and Coordenação de Aperfeiçoamento de Pessoal de Nível Superior (CAPES). All authors acknowledge computational support from the OMICS compute cluster at the University of Lübeck.

## Code availability

No custom algorithms or software have been applied. The R and Python script used for analysis are available from the authors.

## Data availability

Variants rs35044562 and rs67959919 are lead variants of two subsequent Meta-GWAS of the COVID-19 Host Genetics Initiative, both comparing hospitalized COVID-19 with population controls (release 3: ANA_B2_V2 and release 4: B2_ALL, respectively; with summary statistics available at https://www.covid19hg.org). 1000 Genomes variant data (phase 3 release) is available at https://www.internationalgenome.org. Neanderthal variant data is provided by the Max Planck Institute for Evolutionary Anthropology at http://cdna.eva.mpg.de/neandertal (Chagyrskaya, Altai and Vindija 33.19). The world graphic was obtained from Natural Earth, a public domain map dataset.

## Contributions

I.W and V. C.-S conceived the study. All authors designed the study. I.W, V. C.-S. and N.J performed data analysis. I.W. prepared figures and wrote the first manuscript draft. All authors contributed to and approved the final manuscript.

